# A role for prefoldins in H2A.Z deposition in *Arabidopsis*

**DOI:** 10.1101/2021.01.07.425797

**Authors:** Cristina Marí-Carmona, Javier Forment, Miguel A. Blázquez, David Alabadí

## Abstract

The prefoldin complex (PFDc) participates in cellular proteostasis in eukaryotes by acting as cochaperone of the chaperonin CTT. This role is mainly exerted in the cytoplasm where it contributes to the correct folding of client proteins, thus preventing them to form aggregations and cellular damage. Several reports indicate, however, that they also play a role in transcriptional regulation in the nucleus in several model species. In this work, we have investigated how extended is the role of PFDs in nuclear processes by inspecting their interactome and their coexpression networks in yeast, fly, and humans. The analysis indicates that they may perform extensive, conserved functions in nuclear processes. The construction of the predicted interactome for *Arabidopsis* PFDs, based on the ortholog interactions, has allowed us to identify many putative PFD interactors linking them to unanticipated processes, such as chromatin remodeling. Based on this analysis, we have investigated the role of PFDs in H2A.Z deposition through their interaction with the chromatin remodeling complex SWR1c. Our results show that PFDs have a positive effect on SWR1c, which is reflected in defects in H2A.Z deposition in hundreds of genes in seedlings defective in PFD3 and PFD5 activities.

## INTRODUCTION

The prefoldin complex (PFDc) is a heterohexamer present in archaea and eukaryotes, formed by six different subunits named PFD1 to 6 (Liang *et al*., 2020). It was identified as cochaperone of the cytosolic chaperonin CCT, assisting in the folding of actin and tubulins in yeast and in humans (Vainberg *et al*., 1998; Geissler *et al*., 1998). The role of PFDc in tubulin folding seems to be conserved in plants. *Arabidopsis* mutants deficient in individual *PFD* genes have defects associated to impaired tubulin folding, such as hypersensitivity to the microtubule-depolymerizing drug oryzalin or disorganized cortical microtubules (Perea-Resa *et al*., 2017; Rodriguez-Milla and Salinas, 2009; Gu *et al*., 2008). The activity of PFDs in the cytoplasm is not limited to the folding of tubulins and actin, studies in mammals revealed that they exert a prominent role in preventing the formation of protein aggregates, which otherwise compromise cell viability and cause disease (Liang *et al*., 2020). For instance, the Huntingtin protein is overaccumulated in the cytosol of cells knocked down for either *PFD2* or *PFD5,* similar to the overaccumulation of the mutant *huntingtin* protein that causes the Huntington’s disease (Tashiro *et al*., 2013). Recent results have provided molecular insight on the way in which the PFDc prevents protein aggregation and thus contributes to proteostasis (Gestaut *et al*., 2019). By using actin as substrate protein, authors demonstrated that the PFDc remodels actin molecules that are already bound to the CCT but that present a wrong conformation, thus minimizing aggregation and increasing the folding kinetics.

Besides the well-known participation of PFDs in processes that take place in the cytoplasm, an increasing number of reports indicate that they also play roles in the nucleus (Payan-Bravo *et al.*, 2018). These investigations show that PFDs participate in different stages of gene expression, through different mechanisms and with different outcomes. The human PFDN5/MM-1 acts as transcriptional corepressor by recruiting a repressor complex to the c-Myc-bound genomic targets (Satou *et al*., 2001). Although the mechanism is unknown, PFDN1 is recruited to the *Cyclin A* promoter to repress its transcription in humans (Wang *et al.*, 2017). PFD6, on the contrary, acts as transcriptional coactivator through interaction with the transcription factor (TF) FOXO to promote life span in *Caenorhabditis elegans* (Son *et al.*, 2018). PFDs also affect transcription by destabilizing TFs. This is the case of HY5, a TF required for cold acclimation in *Arabidopsis* that is degraded by the 26S proteasome upon interaction with PFD4 (Perea-Resa *et al*., 2017). PFDs also contribute to the activity of the basal transcription machinery. Investigations in the yeast *Saccharomyces cerevisiae* revealed that PFDs bound to the chromatin are required for transcription elongation of long genes by the RNAPII (Millan-Zambrano *et al*., 2013). They also influence post-transcriptional processes. Although through different mechanisms, PFDs affect pre-mRNA splicing both in plants and in humans. Work in *Arabidopsis* has demonstrated that PFDs participate in the splicing of a particular set of pre-mRNAs by promoting the stability of the core spliceosome complex LSM2-8 (Esteve-Bruna *et al*., 2020). In humans, depletion of PFDN5 affects co-transcriptional splicing of long genes by preventing the recruitment of the splicing factor U2AF65 to the target loci (Payán-Bravo *et al*., 2020).

All these results show that PFDs are very versatile proteins, which have the potential to affect the activity of their nuclear protein partners in different ways, ultimately affecting gene expression. Taking into account current evidences and the high degree of conservation of PFDs, we hypothesized that the PFDc or its individual subunits would have a conserved, global role in the regulation of nuclear processes. In this work, we have built a predicted nuclear interactome for *Arabidopsis* PFDs based on the interactors of their orthologs in other model organisms, which has allowed us to identify a role for them in chromatin remodeling.

## RESULTS AND DISCUSION

### Cross-kingdom conservation of interactions between PFD subunits and nuclear proteins

The biological functions of a protein are largely determined by its interactions with other proteins. The first approach to test whether PFDs’ role in the nucleus is general was to perform a data-mining analysis of reported physical interactors of the six PFD canonical subunits, comparing four phylogenetically distant organisms: *S. cerevisiae, Drosophila melanogaster, Homo sapiens,* and *Arabidopsis. S. cerevisiae* canonical PFD subunits have 870 known interactors (Supplementary File 1). Among them, 446 (51.3%) are nuclear, and some are common to 2 or more subunits (Figure S1A). An ontology analysis (Supplementary File 2) revealed a significant enrichment of cytoskeleton components, as expected, but also in chromatin components. Interestingly, the ontology “INO80-type complex” is significantly enriched with 14 proteins, with at least 7 of them being also part of the Swr1 complex (Figure S1A) (March-Diaz and Reyes, 2009). To bolster the idea that these interactions could indicate a functional relationship between PFDs and certain nuclear processes, we examined the genetic interactions described in yeast for the 6 PFD subunits and ranked the results (Figure S1B). Many of the physical interactions with nuclear proteins were further supported by genetic interactions, not only with the interacting PFD subunit, but also with the other ones, suggesting that the nuclear function of the PFDs is probably shared by the entire complex. Again, the interaction with the Swr1 complex was significantly enriched, based on the presence of several Swr1c subunits at the top of the rank. In addition, we searched for the genes that showed statistically significant coexpression with *PFD* genes in yeast, and found that many of them encoded nuclear-localized proteins, with functions related to chromatin remodeling, transcriptional regulation, splicing or DNA recombination and repair (Figure S2). The physical and genetic interactions, together with the significant coexpression values suggest that PFDs may perform extensive nuclear functions in yeast, notably in connection with chromatin remodeling. Indeed, genetic and biochemical studies have demonstrated that several PFDs participate in transcriptional elongation in this model species (Millan-Zambrano *et al*., 2013).

We obtained similar results analyzing fly (Figure S3) and human PFD interactors (Supplementary Files 1): a large group of cytosolic and cytoskeleton proteins and many nuclear interactors involved in splicing, transcription, chromatin and DNA metabolic processes. Interestingly, the functions represented by the human and fly interactors were strikingly conserved, although the identity of the particular interactors was different in most cases (Supplementary File 3). As in the case of yeast, we found two members of the SWR1 complex among the human interactors, but none in *Drosophila.* Available data cover a low percentage of the proteome, specially in *Drosophila,* which may explain these results. Despite the lack of demonstrated physical interaction between PFD and SWR1 c subunits in the fly, we did find significant coexpression between *PFD* genes and the *SWR1c* genes (Figure S4; Supplementary File 4), suggesting that the functional conservation between these two complexes is not only present in fungi, but extends to all metazoans.

The limited protein-protein interaction (PPI) data available for *Arabidopsis* in the BioGRID database shows only 8 PFD interactors. Considering the results obtained for yeast, fly, and humans, we decided to use the predicted interactome for *Arabidopsis,* based on interacting orthologs in several species (Geisler-Lee *et al*., 2007) to gain some insight into the putative conserved roles that PFD could be performing in the nucleus. We used the *Arabidopsis* Interactions Viewer (Geisler-Lee *et al*., 2007) to obtain a network combining the predicted interactome, known PPIs and gene coexpression (Supplementary File 5). As in the case of fungi and animals, the identity of the potential nuclear interactors for the *Arabidopsis* PFD subunits included transcriptional regulators and components of the DNA recombination and repair machinery, as well as proteins involved in RNA splicing (Figure 1). Noteworthy, a large set of interactors were involved in chromatin remodeling, including subunits of the SWR1c or SWI/SNF complexes.

**Figure 1.**
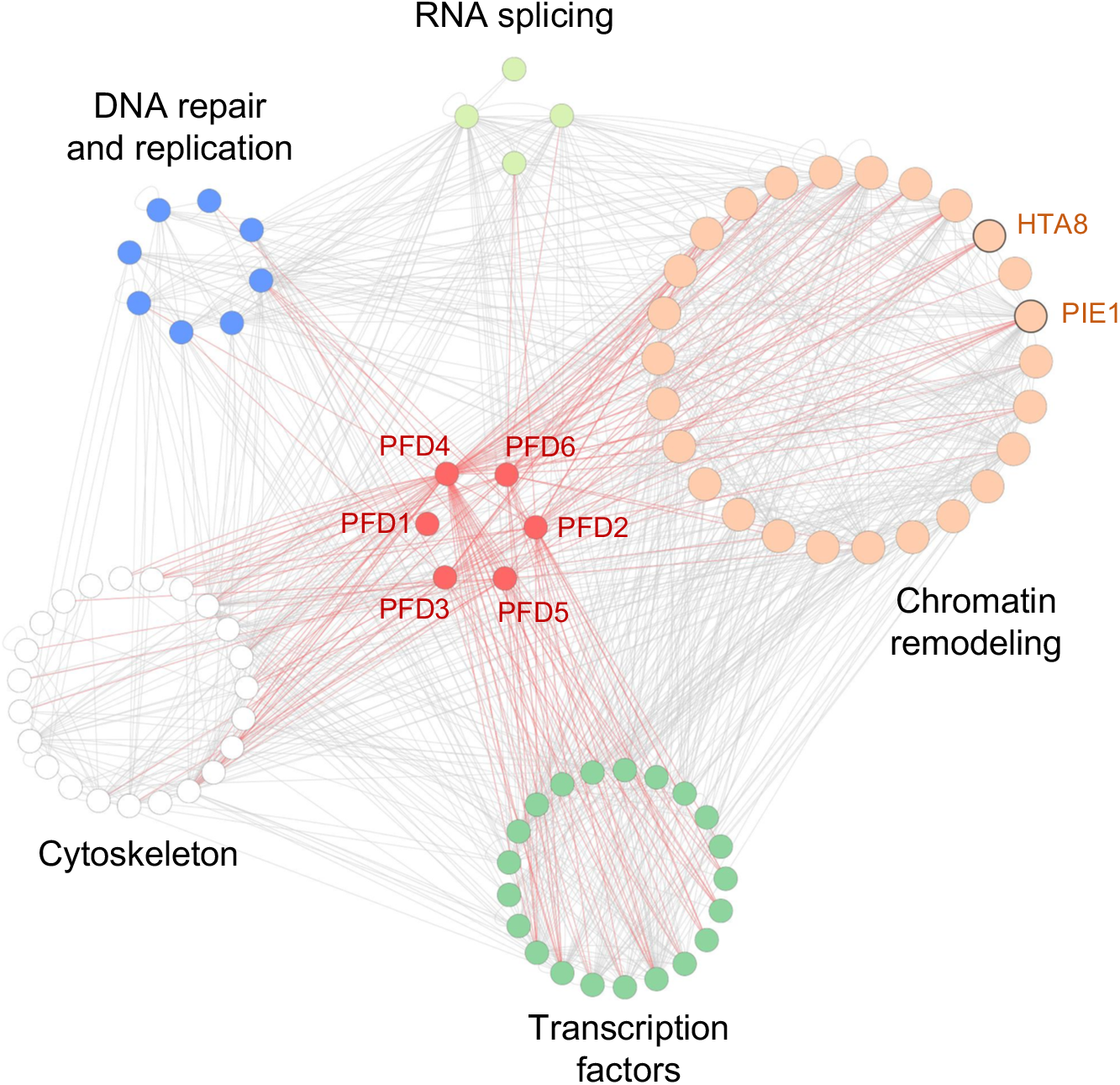
The predicted nuclear interactome of PFD in *Arabidopsis.* Physical interactions and co-expression values were obtained from the *Arabidopsis* Interaction Viewer. Red edges represent statistically significant co-expression of genes with nuclear localization and PFD subunits.

Given the evidence for an extensive role of PFD in the nucleus, we set out to confirm the physical and functional connection between these proteins and at least one of the nuclear complexes identified in the *in silico* analyses. We decided to focus on the SWR1 chromatin remodeling complex because it is specially represented, with numerous subunits present in our interaction data for fly, humans, yeast, and the predicted *Arabidopsis* network.

### PFD interacts physically with SWR1c in *Arabidopsis*

We performed a targeted yeast two-hybrid screening to test the interactions between *Arabidopsis* PFD and most of the SWR1c subunits (March-Diaz and Reyes, 2009). We found that each PFD subunit interacted with at least one of the components of SWR1 c, but not with the three H2A.Z histones (Figure 2A). This result is compatible with a model in which the interaction with SWR1c is established by the PFD complex, rather than by individual subunits. Although the structure of the plant SWR1c has not been elucidated, it is possible to use the yeast structure as a likely model (Nguyen *et al*., 2013). Besides the SWR1 ATPase, three subcomplexes can be identified, the N-module, the C-module, and the RVB1/2 module. Considering the individual interactions detected by yeast two-hybrid, we hypothesized that the interaction surface with the PFDc would be mapped to the face opposite to the SWR1 ATPase with which the nucleosome interacts.

**Figure 2.**
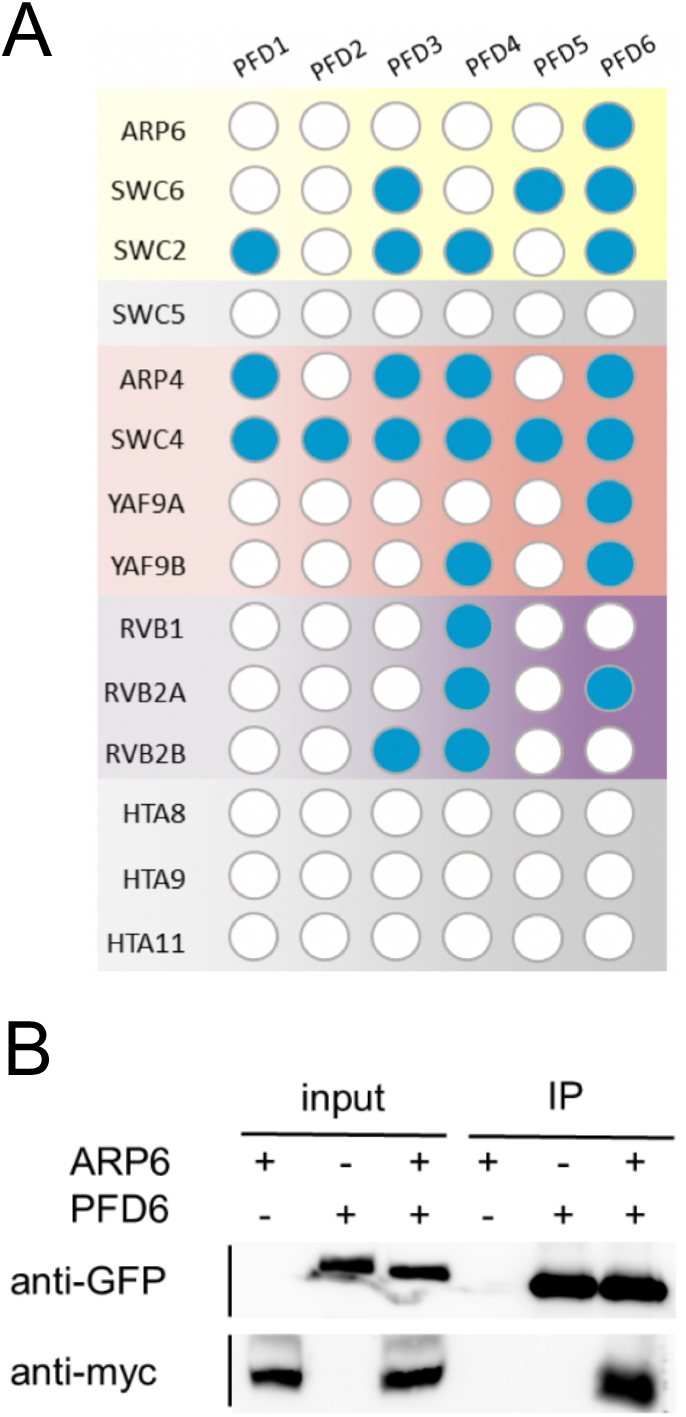
Physical interaction between PFD and the SWR1 complex. (A) Results of the yeast two-hybrid assays. Blue and white circles indicate interaction or no interaction, respectively. (B) Interaction between cMyc-ARP6 and YFP-PFD6 detected by coimmunoprecipitation in agroinfiltrated leaves of *N. benthamiana*.

Interestingly, affinity purification followed by mass spectrometry has also identified PFDs as ARP6- and SWC4-interacting proteins (Gomez-Zambrano *et al*., 2018; Luo *et al*., 2021; Potok *et al*., 2019; Sijacic *et al*., 2019). To obtain additional proof for the *in vivo* interaction between PFD and SWR1 subunits, we performed a co-immunoprecipitation assay between cMyc-ARP6 and YFP-PFD6 expressed in *Nicotiana benthamiana* leaves (Figure 2B). We detected cMyc-ARP6 after immunoprecipitation with anti-GFP only in leaves co-expressing both fusion proteins, confirming that the two proteins interact physically in plant cells.

### PFDs contribute to the flowering time by affecting H2A.Z levels in the *FLC* locus

To explore if the interaction is physiologically relevant, we decided to investigate the flowering-time phenotype of *pfd* mutants, given that the SWR1c has been involved in the regulation of this process (Lazaro *et al*., 2008; Martin-Trillo *et al*., 2006). It has also been described that they act in the ambient temperature pathway for flowering (Kumar *et al*., 2012), so we decided to analyze the flowering time of *arp6, pfd3,* and *pfd5* mutants at 16, 22, and 27° C under short days (SD, Figure 3A), because the effect of the thermal induction of flowering is more marked in this condition. Consistent with previous reports, *arp6* flowered earlier than wild-type plants at all temperatures. At 22° and 16° C, *pfd* mutants showed early flowering, but at 27° C there was no significant difference with the wild-type. The flowering-time defect of *pfd* mutants was less pronounced than that of *arp6* but showed the same tendency. The *swc6* mutants also displayed early flowering at low temperature, and there was only a marginal increase in this defect when combined with the *pfd3* mutation, being the phenotype similar to the *h2a.z* mutant that is highly deficient in H2A.Z levels (Figure 3B) (Coleman-Derr and Zilberman, 2012). These results would be consistent with PFDs and SWR1 c mostly acting together, at least for the control of flowering time.

**Figure 3.**
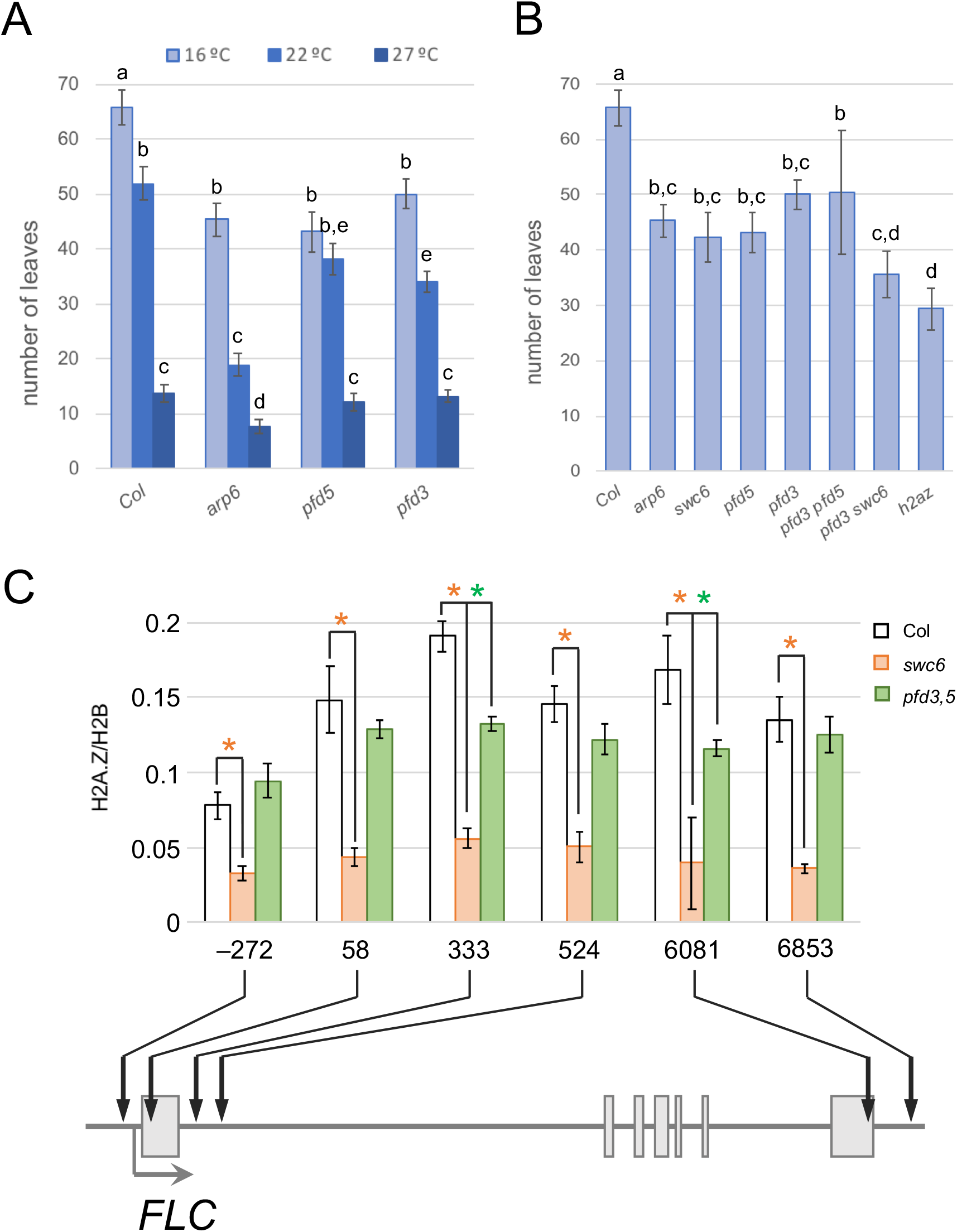
Flowering-time defects of *pfd* mutants. (A) Total number of leaves produced by wild-type, *arp6*, *pfd3*, and *pfd5* mutants before flowering at three different growth temperatures under short days. (B) Total number of leaves of plants with *pfd* and *swr1c* mutant genotypes at 16°C under short days. Small-caps letters indicate similar groups in an ANOVA test (n≥12, p<0.01). (C) H2A.Z relative levels in the *FLC* locus. Plants were grown for 14 days in SD, just before the onset of flower formation in the earlier-flowering genotype, and H2A.Z levels were determined by chromatin immunoprecipitation (ChIP) coupled to qPCR using primers designed for the different regions of the *FLC* gene, indicated by arrows. Numbers indicate the position relative to the Transcription Start Site (TSS). Error bars indicate standard error (n=3). Asterisks mark statistically significant differences with respect to the wild-type values (Student’s t-test, p<0.05).

The early flowering phenotype of mutants in the SWR1 c can be explained by the reduced deposition of H2A.Z over the locus of the flowering repressor *FLC* that results in reduced expression (Deal *et al*., 2007). Given the early flowering phenotype of the *pfd* mutants and the physical interaction between PFD and SWR1c, we wondered whether this phenotype could be caused by reduced deposition of H2A.Z in the *FLC* locus as well.

For that purpose, we analyzed the distribution of H2A.Z in *FLC* by ChIP-qPCR in the wild-type and in the *pfd3,5* and *swc6* mutants grown for 14 days under SD conditions. Results in Figure 3C show that H2A.Z accumulated around the 5’ and 3’ ends of the gene in the wild-type, as described (Deal *et al*., 2007). The *swc6* mutation strongly affected the accumulation of the histone variant, being reduced over the whole gene. This result is similar to the effects reported for mutants in other subunits (Deal *et al.*, 2007). Importantly, we also observed a reduction in H2A.Z deposition in the *pfd3,5* mutant, being the levels intermediate between the wild-type and the *swc6* mutant.

These results not only suggest that PFD activity may contribute to the control of flowering time through SWR1c, highlighting the physiological relevance of the interaction between both complexes, but, more importantly, they indicate that PFDs may have a positive role in SWR1 c activity. Given these results, we decided to study the impact of the interaction at genomic level.

### Transcriptomic analysis of PFDc and SWR1c loss-of-function mutants underscores overlapping functions

To obtain a more general picture of the effect of PFDs upon SWR1c, we examined the genome-wide transcriptional defects of *pfd* mutants and compared them with those caused by loss of SWR1c activity. To this end, we performed an RNA-seq analysis on the *pfd3,5, arp6,* and *swc6* mutants. All plants were grown at 22° C in SD conditions, and samples were collected 2 weeks after germination. Our analysis identified around 2700 misregulated genes (fold change [FC] ≥ 1.5; false discovery rate [FDR] ≤ 0.05) in *arp6* and *swc6*, with half of them being up-regulated and half of them down-regulated. Moreover, a vast majority (70%) were misregulated in both mutants (Figure 4A). An equivalent analysis of *pfd3,5* yielded 368 genes up-regulated and 798 genes down-regulated (Figure 4A). 46.7% of misregulated genes in *pfd3,5* overlapped with misregulated genes in *arp6* and *swc6*. Importantly, the transcriptional changes in the *pfd3,5* mutant followed the same tendency as the changes in the mutants affecting the SWR1c (Figure 4A). This can be also observed in the heatmap of Figure 4B, where the fold change of every misregulated gene common to all mutants is represented in a color scale, and genes are grouped in four clusters according to their behavior.

**Figure 4.**
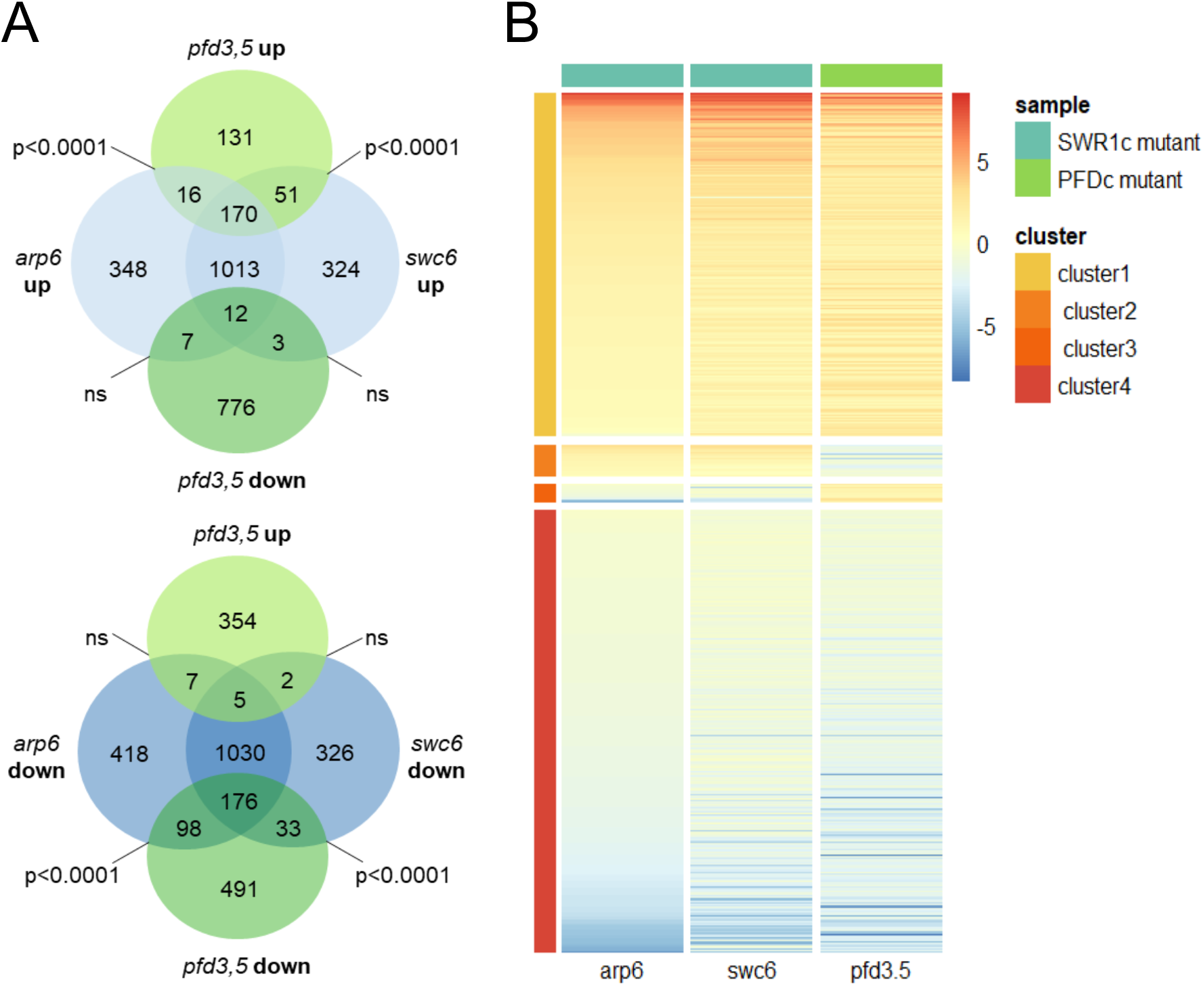
Transcriptomic analysis of the *pfd3,5, swc6,* and *arp6* mutants. (A) Venn diagrams showing the number of genes misregulated in each mutant (FC≥1.5, FDR≤0.05) and the overlap between them. Statistical significance of the overlap was calculated with a Fisher’s exacts t-test. (B) Heatmap highlighting the overlap in the sense of transcriptional changes in the PFDc and SWR1c loss-of-function mutants. Only the genes jointly misregulated in the three genotypes are shown.

Gene Ontology over-representation analysis among the genes misregulated in the two SWR1c mutants showed an enrichment in biological processes mostly related to defense response, response to stimulus and cell death, and also in categories like ‘post-embryonic plant development’ and ‘mitotic cell cycle’ (Figure S5; check Supplementary File 6 for a detailed list). These results are consistent with previous reports (Berriri *et al*., 2016; Dai *et al*., 2017; Sura *et al*., 2017). Importantly, many of these functions were also enriched among the genes misregulated in the *pfd3,5* mutant (Figure S5 and Supplementary File 6). The overlap in the gene targets and biological processes affected by mutants in the two complexes indicate that the collaboration of the PFDc with SWR1 c is rather extensive and not restricted to the regulation of H2A.Z deposition at the *FLC* locus.

### PFD affects H2A.Z deposition in a subset of genes

To further extend the study of the functional relationship between PFDs and SWR1c, we analyzed the genome-wide distribution of the histone H2A.Z in the wild-type and in *pfd3,5* and *swc6* mutants by performing a ChIP-seq experiment on 14-day old seedlings grown at 22° C in SD conditions. The two biological replicates were highly similar, so we decided to pool them for further analysis. After peak calling, 18841 and 11320 peaks were detected in the wild-type and *swc6* seedlings (Supplementary File 7), respectively, localized mainly in genic features (Figure 5A). The accumulation profile of H2A.Z in the wild-type showed strong enrichment around the transcription start site (TSS) (Figure 5B and C) (Yelagandula *et al*., 2014; Coleman-Derr and Zilberman, 2012; Sura *et al*., 2017; Gomez-Zambrano *et al*., 2018), while the same distribution but with strongly decreased levels were observed in the *swc6* mutant, as reported, for instance, for the *swc4* mutant (Gomez-Zambrano *et al*., 2018).

**Figure 5.**
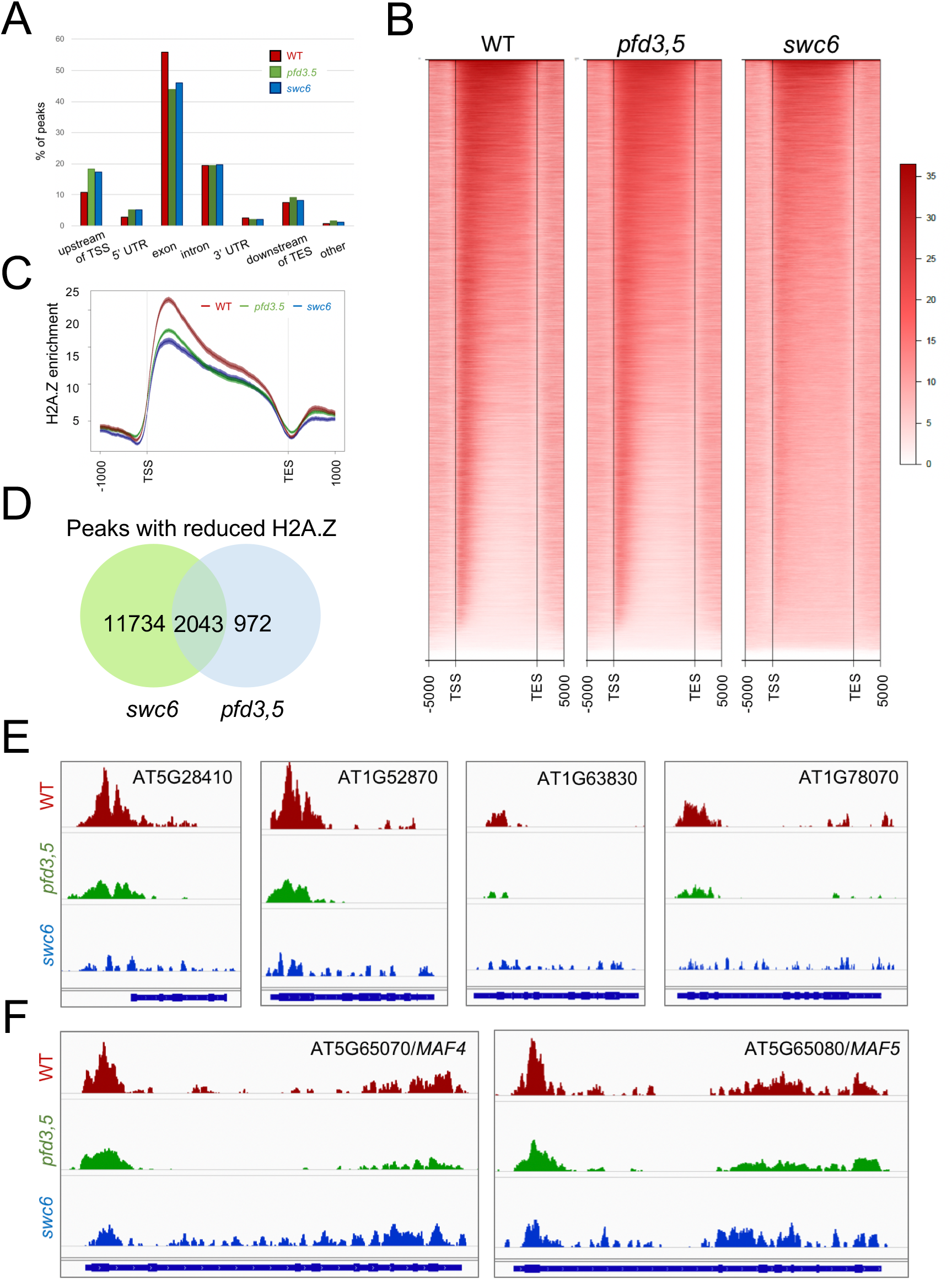
Genomic distribution of H2A.Z in wild-type, *pfd3,5*, and *swc6* seedlings. (A) Distribution of H2A.Z in the different gene features. (B) Heatmap showing the H2A.Z enrichment in the genes occupied in the wild-type and ranked from top to bottom according to the enrichment in the wild-type. (C) Metagene plot of average H2A.Z enrichment in the three genotypes of the gene set affected in *pfd3,5* mutant seedlings. (D) Venn diagram showing the overlap in peaks with reduced H2A.Z deposition. TSS transcription start site, TES transcription stop site. (E) IGV view of the H2A.Z occupancy in four representative genes. (F) IGV view of the H2A.Z occupancy in the *FLC* paralogs *MAF4* and *MAF5.* All tracks are equally scaled. The exon/intron structures of the corresponding genes are shown at the bottom of each panel.

19116 peaks were detected in *pfd3,5* seedlings (Supplementary File 7), with slight reduction in the peak around the TSS in this mutant compared to the wild-type (Figure 5B). Indeed, a statistical analysis identified 3156 peaks with modest but significant decrease in H2A.Z enrichment in the *pfd3,5* mutant (p adj. <0.01, Supplementary File 8). We also detected 628 peaks with increased enrichment. A metagene plot representing only the subset of genes with reduced enrichment in the *pfd3,5* mutant clearly shows the reduction in the H2A.Z levels (Figure 5C). Importantly, 63.4% of the of the peaks with decreased H2A.Z in *pfd3,5* (2627 peaks) showed also decreased levels in *swc6* plants (Figure 5D). For instance, see examples of the H2A.Z distribution in four top ranked genes affected in the *pfd3,5* mutant (Figure 5E). Although the reduction of H2A.Z in *FLC* was not statistically significant in our analysis, a clear reduction over the paralog gene loci *MAF4* and *MAF5* was observed in the *pfd3,5* seedlings (Figure 5F), suggesting that misregulation of these two genes may also contribute to the early-flowering phenotype of *pfd* mutants (Figure 3).

The distribution of H2A.Z along one gene depends on its transcriptional behavior. We wondered if PFDs affects H2A.Z deposition depending on the transcription level of the gene. For that purpose, we first grouped genes in six groups of expression based on the expression level found in our RNA-seq analysis of wild-type seedlings, and then plotted the average H2A.Z levels for the wild-type, *pfd3,5* and *swc6* in each group (Figure S6). Results showed that H2A.Z deposition was affected in the *swc6* mutants regardless of the expression level of the gene, as expected for a mutant affecting a core subunit of the SWR1 c. Nonetheless, defects were most apparent in gene groups with lower expression levels in the *pfd3,5.* Note that the H2A.Z distribution in the gene group with the highest expression level in the *pfd3,5* mutant is indistinguishable of the wild-type. These results suggest that PFDs’ contribution to H2A.Z deposition is more relevant in genes with lower expression level, likely associated to gene responsiveness. This agrees with the Gene Ontology analysis (Figure S7) that showed that several categories associated to responses to the environment were enriched among the common genes affected in *swc6* and *pfd3,5* mutants.

### Network analysis identifies candidate TFs acting downstream of PFD-SWR1c

In agreement with previous observations (Gomez-Zambrano *et al*., 2018) in which most genes downregulated in the *swc4* mutant did not suffer from defects in H2A.Z deposition, we did not find a large overlap between the sets of misregulated genes and those with reduced H2A.Z in *pfd3,5* (Figure 6A). Although this overlap was statistically significant, only 4% of the genes with defective H2A.Z deposition were downregulated in the case of *pfd3,5,* and 5% in the case of *swc6* (Figure 6A). Thus, it is likely that most of the genes misregulated in *pfd3,5* are indeed targets of only a handful of TFs whose expression is regulated by H2A.Z deposition. To investigate this possibility and identify the primary targets for PFD-dependent SWR1c activity, we extracted the list of TFs present among the 798 downregulated genes in *pfd3,5* which had also been tagged for defective H2A.Z levels. We found only 7 matching these criteria (Figures 6B and C). To find the possible connections between these 7 TFs and the rest of the genes downregulated in *pfd3,5,* we used the TF2Network tool (Kulkarni *et al*., 2018). This algorithm identifies putative regulatory relationships based on experimental genome-wide evidence of TF binding to promoters, and co-expression values. With those criteria, we found that the first tier of 7 TFs were predicted to regulate 752 out of the 798 downregulated genes (p<0.01 as confidence threshold value) (Figure 6D, Supplementary File 9). Moreover, this set of 752 genes formed a very robust network because the algorithm also highlighted the enrichment of a second tier of 8 additional TFs whose H2A.Z levels were not significantly affected in the *pfd3,5* mutant, but their expression levels were lower and were direct targets of the first-tier TFs (Figure 6D, Supplementary File 9).

**Figure 6.**
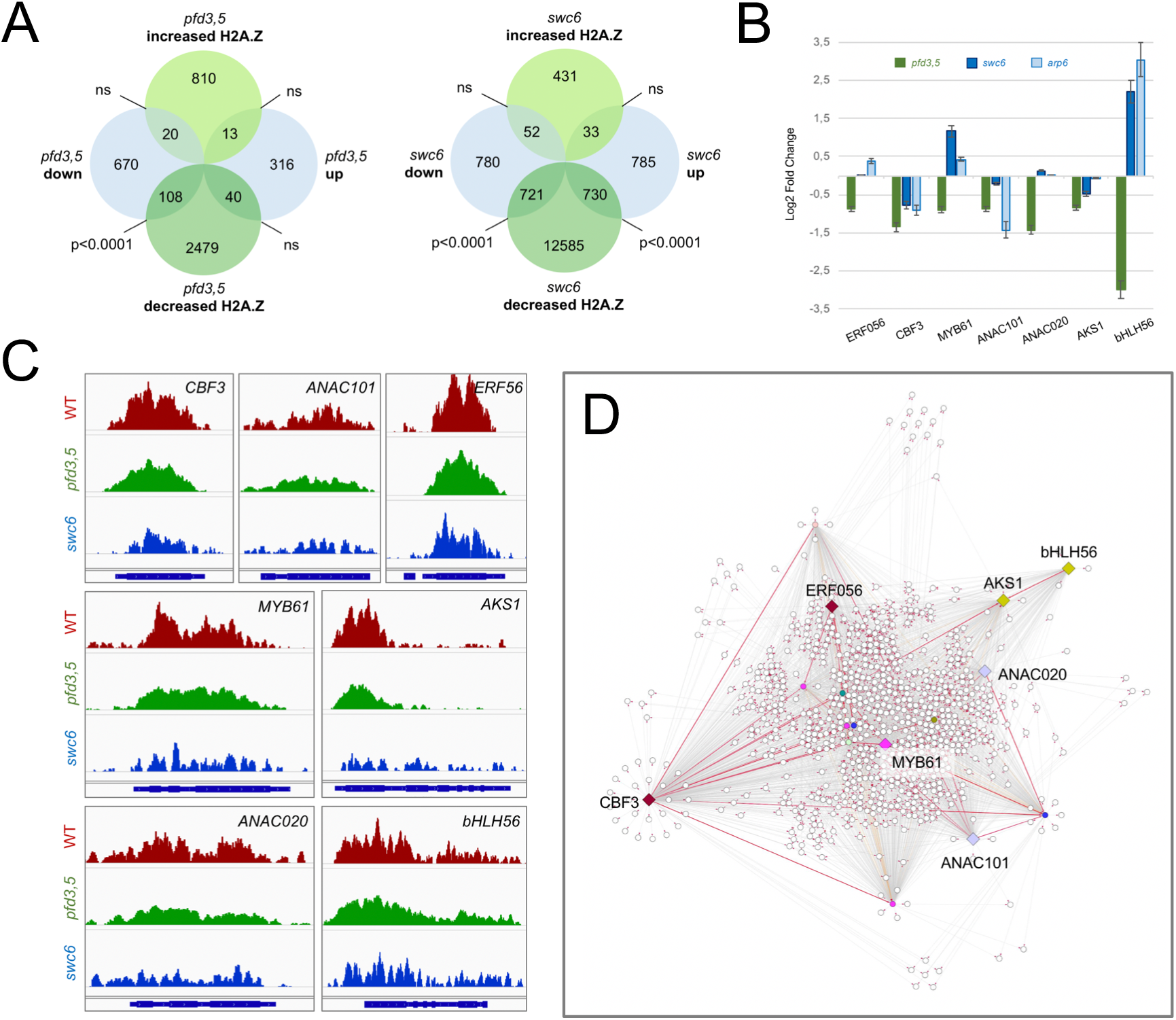
(A) Overlap between genes with differential H2A.Z accumulation and misexpressed genes in *pfd3,5* and *swc6* mutants. (B) Expression levels of selected TFs with defective H2A.Z deposition. The Log2 Fold Change values were extracted from the RNA-seq experiment. (C) IGV view of H2A.Z distribution in the 7 selected TFs that display lower expression levels in the *pfd3,5* mutant. All tracks are equally scaled. The exon/intron structures of the corresponding genes are shown at the bottom of each panel. (D) Hierarchical interaction analysis of the coregulation of the genes downregulated in *pfd3,5* mutants by TFs affected in H2A.Z deposition in the same mutant. Colored nodes indicate TFs in the set of downregulated genes which are predicted to regulate other genes in the same set. Diamonds mark those TFs that also show defects in H2A.Z deposition in the *pfd3,5* mutant. Red edges highlight transcriptional regulation between TFs. Red arrowheads indicate the direction of the regulation. Hierarchical interactions were calculated and represented with Cytoscape.

These results provide an explanation for the large number of genes whose expression is regulated by PFDs but do not seem to have H2A.Z defects in *pfd* loss-of-function mutants. However, they do not explain two observations: why many of the genes with altered H2A.Z deposition in *pfd* or *swc6* do not have defects in expression levels, and why only a small subset of loci regulated by SWR1c are also regulated by PFDs. To clarify this second issue, we decided to investigate the possible molecular mechanisms by which PFD could affect SWR1c activity.

### Possible molecular mechanisms of PFD effect on SWR1c

Given the physical interaction found between PFD6 and ARP6, and the role of PFD as a co-chaperone, we considered that a plausible possibility is that PFDs directly alter SWR1c activity by affecting ARP6 protein levels. We analyzed ARP6 protein levels in *swc6*, *pfd3,* and *pfd5* mutant backgrounds (Figure S8A). We found lower levels of ARP6 in the *swc6* mutant compared to the wild-type. This result suggests that loss of one core subunit of the complex leads to lower accumulation of another, likely as consequence of decreased stability when the whole complex cannot be assembled. Importantly, no apparent changes in the level of ARP6 was found in any *pfd* mutant. This analysis indicates that impairment of PFD function has no impact on ARP6 protein levels.

According to the role of PFD as co-chaperones of the chaperonin CCT, recent results indicate that the PFDc assistance to CCT is also required in the nucleus of human cells, in particular, for the assembly of the histone deacetylase HDAC1 into transcriptional repressor complexes (Banks *et al*., 2018). Thus, we next tested the possibility that PFDs are required for the assembly or for keeping the integrity of SWR1c. To test this, we subjected extracts of WT and *pfd3,5* seedings to size exclusion chromatography. The SWR1 c eluted in fractions corresponding to the reported size of the complex (Deal *et al*., 2007) (Figure S8B). The elution profile was very similar in the *pfd3,5* mutant. This result indicates that the lack of at least PFD3 and PFD5 activities does not have any effect in the assembly or integrity of the SWR1 c.

We next investigated if PFDs affect the recruitment of the SWR1c to the chromatin. For that purpose, we performed a cellular fractionation of extracts of seedlings of *pfd3, pfd3,5,* and *swc6* mutants and followed the presence of the complex in each fraction by immunoblots with anti-ARP6 antibodies. We found the ARP6 protein in all fractions (Figure S8C). This result was surprising, as we expected it to be exclusively in nuclear fractions. Nonetheless, its presence in the cytoplasm may suggest that the complex is assembled in this location, prior to its import into the nucleus. Interestingly, we found a higher proportion of ARP6 in the nucleoplasm of *pfd3* and *pfd3,5* mutants than in the wild-type, although no apparent differences were observed in the chromatin. The accumulation of ARP6 in the nucleoplasm of the mutants may have a small effect, yet not noticeable by western analysis, in the pool of chromatin-bound SWR1c, affecting only H2A.Z deposition in loci highly sensitive to SWR1c levels. This possibility is in agreement with the ChIP-seq data, which indicate that PFD’s effect on H2A.Z deposition is restricted to a subset of genes.

## CONCLUDING REMARKS

Our work shows that the function of PFDs in nuclear processes may be quite extended, a view that could not be anticipated after the discovery of these proteins acting as cochaperones in the cytosol. The comparative analysis of the PFD interactome and coexpression network in various model organisms indicates that many of the functions that PFDs exert in the nucleus are conserved, suggesting that those functions were fixed early during evolution. Despite of this, species specific functions also arose. For instance, the nuclear presence of PFDs in the nucleus in *Arabidopsis* is promoted by the plant-specific transcription regulators DELLA proteins (Locascio *et al*., 2013; Perea-Resa *et al*., 2017).

The comparative analysis has allowed us to define a novel role for PFDs in chromatin remodeling. Although our results do not provide a molecular mechanism to explain how PFDs promote the activity of the chromatin remodeler SWR1 c, genomic analyses clearly show that PFDs are required for the proper deposition of the H2A.Z variant in a set of gene loci. The fact that the effect of *pfd* mutations is not observed over all loci normally occupied by H2A.Z, suggests to us that the effect of PFDs on SWR1c may occur locally at the chromatin of the target genes. This may occur, for instance, by facilitating the recruitment of the remodeler to the chromatin. Although PFDs do not seem to bind DNA directly, they have been found bound to the chromatin by ChIP in yeast and humans (Wang *et al*., 2017; Payán-Bravo *et al*., 2020; Millan-Zambrano *et al*., 2013), and by cell fractioning in *Arabidopsis* (Locascio *et al*., 2013). The molecular mechanism that provides specificity to PFDs’ action on H2A.Z deposition is, therefore, a matter for future research.

## MATERIALS AND METHODS

### *In silico* analysis

Protein-protein and genetic interaction data were retrieved from BioGrid (https://thebiogrid.org/), IntAct (https://www.ebi.ac.uk/intact/), and MINT (https://mint.bio.uniroma2.it/) databases. Coexpression and Gene Ontology (GO) annotation data were retrieved from InterMine (http://intermine.org/) and YeastNet v3 (https://www.inetbio.org/yeastnet/). Additionally, a list of yeast chaperone interactors was obtained from (Gong *et al.*, 2009). The *Arabidopsis* predicted interactome was obtained from (Geisler-Lee *et al.*, 2007). Networks were depicted with Cytoscape software (https://cytoscape.org/).

### Plant material and growth conditions

All plant lines were in Col-0 background. We used previously characterized T-DNA insertion lines: *arp6-1, swc6-1, h2a.z, pfd3,* and *pfd5.* The *pfd3,5* and *pfd3 swc6-1* mutants were generated by genetic crosses of the respective single mutant lines. Seeds were sown on ½ MS medium (Duchefa), 0.8% (w/v) agar (pH 5.7) and stratified at 4 °C in the dark for 4 days. Unless otherwise stated, plants were grown on MS plates for 2 weeks under short-day photoperiod (8h light/16 h dark), with fluorescent white light intensity of 100 μmol m^-2^ s^-1^, at 22 °C.

For flowering time analysis, plants were germinated on MS plates at 22 °C under short-day conditions. After 7 days, seedlings were transferred to soil and moved into growth chambers at 16, 22 or 27 °C. Flowering time was determined by counting the total leaf number after the first flower opened.

### Yeast two-hybrid assays

The coding sequence (CDS) of PFD subunits, SWR1c subunits, HTA8, HTA9, and HTA11 were transferred to both pGADT7 and pGBKT7 plasmids (Clontech) by Gateway technology. Haploid yeast strains Y2HGold and Y187 (Clontech) were transformed with pGBKT7 and pGADT7 constructs, respectively. Diploid cells carrying both plasmids were generated by mating and interaction assays were performed on synthetic complete minimal medium lacking His, Leu, and Trp, and supplemented with 0-5 mM of 3-amino-1,2,4-triazole (3-AT).

### Protein co-immunoprecipitation assay

The CDSs of PFD6 and ARP6 were transferred to the pEarleyGate104 and pEarleyGate203 vectors (Earley *et al.*, 2006), respectively, to generate the fusions YFP-PFD6 and Myc-ARP6. Constructs were introduced into *Agrobacterium tumefaciens* C58 cells, which were used to infiltrate *Nicotiana benthamiana* leaves, in combination and individually. Leaf samples were collected after 3 days. Frozen ground tissue was homogenized with extraction buffer (50mM Tris-HCl pH 7.0, 10% glycerol, 1mM EDTA pH 8.0, 150 mM NaCl, 10mM DTT, and 1X protease inhibitor cocktail [cOmplete EDTA free, Roche]) in a ratio 2:1 (v/v), and incubated on ice for 15 min. Extracts were centrifuged twice for 10 min at 14000 rpm at 4 °C, and proteins were quantified by Bradford assay. 50 μg of total proteins were denatured in 1X SDS buffer and used as input sample. 1.5 mg of total proteins were incubated with 50 μL of anti-GFP paramagnetic beads (Miltenyi) for 2 h at 4 °C in a rotating wheel, and then loaded onto μcolumns (Miltenyi). Columns were washed with 800 μL of cold extraction buffer and proteins were eluted with 1X SDS buffer, following manufacturer’s instructions. Input and immunoprecipitated samples were analyzed by Western blot.

### Size exclusion chromatography

Extracts of wild-type and *pfd3,5* seedlings were prepared in extraction buffer (50 mM Tris-HCl pH 7.5, 150 mM NaCl, 10 mM MgCl2, 10% Glycerol, 0.5% Nonidet P-40, 2 mM PMSF and 1X protease-inhibitor cocktail). Proteins were loaded in a SuperoseTM 6 Increase (GE Healthcare) column. Fourteen fractions (250 μL each) were collected and those where ARP6 is present are shown. Proteins in fractions were precipitated in 10% trichloroacetic acid on ice for 90 min and then washed twice with cold acetone before Western-blot analysis.

### Subcellular fractionation

Subcellular fractionation was performed according to (Zhang *et al*., 2014), with minor modifications. 1.5 grams of seedlings were ground in liquid nitrogen and homogenized in 3 mL of cold Honda buffer (0.44 M Sucrose, 20 mM HEPES KOH pH 7.4, 2.5% Percoll, 5% Dextran T40, 10 mM MgCl2, 0.5% Triton X-100, 5mM DTT, 1 mM PMSF, and 1x protease inhibitor cocktail). The homogenate was filtered through two layers of Miracloth and centrifuged at 2000 g at 4 °C for 5 min. 1 mL of the supernatant was centrifuged at 10000g at 4 °C for 10 min, and the supernatant collected as cytoplasmic fraction. The first pellet was resuspended in 1 mL of Honda buffer and centrifuged at 1800g for 5 min to pellet the nuclei. The pellet was washed 4 times with Honda buffer, rinsed with 1X PBS buffer (137 mM NaCl, 2.7 mM KCl, 10 mM Na2HPO4, 2 mM KH2PO4) with 1mM EDTA, and resuspended with 150 μL of cold glycerol buffer (20 mM Tris-HCl pH 7.9, 50% glycerol, 75 mM NaCl, 0.5 mM EDTA, 0.85 mM DTT, 0.125 mM PMSF, and 1x protease inhibitor cocktail), to which 150 μL of cold nuclei lysis buffer was added (10 mM HEPES KOH pH 7.4, 7.5 mM MgCl2, 0.2 mM EDTA, 0.3 M NaCl, 1 M urea, 1% NP-40, 1 mM DTT, 0.5 mM PMSF, 10 mM β-mercaptoethanol, and 1x protease inhibitor cocktail). This was vortexed twice for 2 s and incubated on ice for 2 min, following centrifugation at 14000 rpm at 4 °C for 2 min. The supernatant was collected as nucleoplasmic fraction. The chromatin pellet was rinsed with 1X PBS/1mM EDTA and resuspended in 150 μL cold glycerol buffer and 150 μL cold nuclei lysis buffer. Protein concentrations were determined by using the Pierce 660 nm protein assay (ThermoFisher Scientific) according to the manufacturer’s instructions. The fractions were analyzed by Western blot.

### Immunoblots

Protein samples were separated by SDS-PAGE, transferred to PVDF membranes, and immunolabeled with specific antibodies against anti-c-Myc (1:1000, E910, Roche), anti-GFP (1:10000, Living Colors, JL-8), anti-ARP6 (1:200, Kerafast, EGA929), anti-DET3 (1:10000, provided by Karin Schumacher), or anti-H3 (1:5000, Abcam, ab1791).

Horseradish peroxidase-conjugated anti-rabbit (Agrisera) and anti-mouse (Agrisera) were used as secondary antibodies at 1/20,000 and 1/10,000 dilutions, respectively. Chemiluminiscence detection was performed with the Supersignal West FEMTO maximum sensitivity substrate (Thermo-Fisher Scientific) and protein bands were detected using the LAS-3000 Imaging system (Fujifilm).

### RNA-seq and RNA-seq data analysis

Total RNA was extracted from Col-0, *arp6-1, swc6-1* and *pfd3,5* seedlings (three biological replicates) using RNeasy Plant Mini Kit (Qiagen) according to the manufacturer’s instructions. The RNA concentration and integrity [RNA integrity number (RIN)] were measured in a RNA nanochip (Bioanalyzer, Agilent Technologies 2100) by the IBMCP Genomics Service. Library preparation and sequencing were performed by the Genomics Service of the University of Valencia. Reads were mapped to the *Arabidopsis* TAIR10 reference genome using Bowtie 2 (Langmead and Salzberg, 2012), and counts were calculated with HTSeq-count software (Anders *et al*., 2015) and differentially expressed genes (DEG) were identified with edgeR Bioconductor package (McCarthy *et al*., 2012). DEGs were selected according to a fold change cut-off >|1.5| and a p value <0.05. Normalized expression values for each gene were calculated as transcripts per kilobase million (TPM). Heatmap representation was generated with Pheatmap R package. Gene Ontology annotation and over-representation of biological processes terms were performed with the clusterProfiler Bioconductor package (Yu *et al.*, 2012) using p-value and q-value cut-offs of 0.01. Redundancy of enriched Gene Ontology terms was reduced and results were represented in a figure using GOSemSim Bioconductor package (Yu *et al.*, 2010), with similarity cut-off of 0.7 on level 3 Gene Ontology terms.

### ChIP experiments

Chromatin was extracted and immunoprecipitated from Col-0, *swc6-1* and *pfd3,5* seedlings (two biological replicates) as described in (Gallego-Bartolomé *et al*., 2019), using anti-HTA9 (10 μg, Agrisera, AS10 718) and anti-H2B (3μg, Abcam, ab1790). For ChIP-seq, library preparation and sequencing were carried out by the CRG Genomics Core Facility (CRG, Barcelona, Spain). For ChIP-qPCR, amplification was performed using 7500 Fast Real-Time PCR System (Applied Biosystems) with SYBR Premix Ex Taq II (Tli RNaseH Plus) ROX plus (Takara Bio), with primers previously described in (Yang *et al*., 2014). Two biological replicates, each one including three technical qPCR replicates, were performed. Results are given as the percentage of input normalized to histone H2B.

### ChIP-seq data analysis

Read mapping to the TAIR10 reference genome was performed using Bowtie 2. Intersection of peaks called in independent biological replicates confirmed the consistency between both replicates, which allowed us to pool them. Peak calling and differential enrichment analysis were carried out with the software SICER 2 (Zang *et al*., 2009), using input as the control library with a redundancy threshold of 1, a window size of 200 bp, a gap size of 600 bp, and FDR = 0.01. The heatmaps and metagene plots were generated with SeqPlots (Stempor and Ahringer, 2016) using the normalized coverage files generated by SICER 2. Annotation of peak location relative to different genomic features was performed using PAVIS (Huang *et al*., 2013) with default parameters.

### RNA-seq and ChIP-seq meta-analysis

For the meta-analysis of gene expression levels and H2A.Z enrichment, genes were split into 6 groups based on TPM in wild-type seedlings (TPMM<1, TPM=1-5, TPM=5-10, TPM=10-30, TPM = 30-150, TPM>150), and mean H2A.Z enrichment was represented for each group with SeqPlots (Stempor and Ahringer, 2016). TF2Network tool (Kulkarni *et al.*, 2018) was used to find over-represented TFs binding sites, and this information together with expression levels and H2A.Z enrichment levels were used to build a hierarchical network using Cytoscape software (https://cytoscape.org/).

## ACKNOWLEDGEMENTS

This work has been funded by grants from the Agencia Española de Investigación BIO2013-43184-P to D.A. and M.A.B. and BIO2016-79133-P to D.A. C.M-C. was recipient of a predoctoral fellowship from the Universidad Politécnica de Valencia.

---

**Figure S1.**
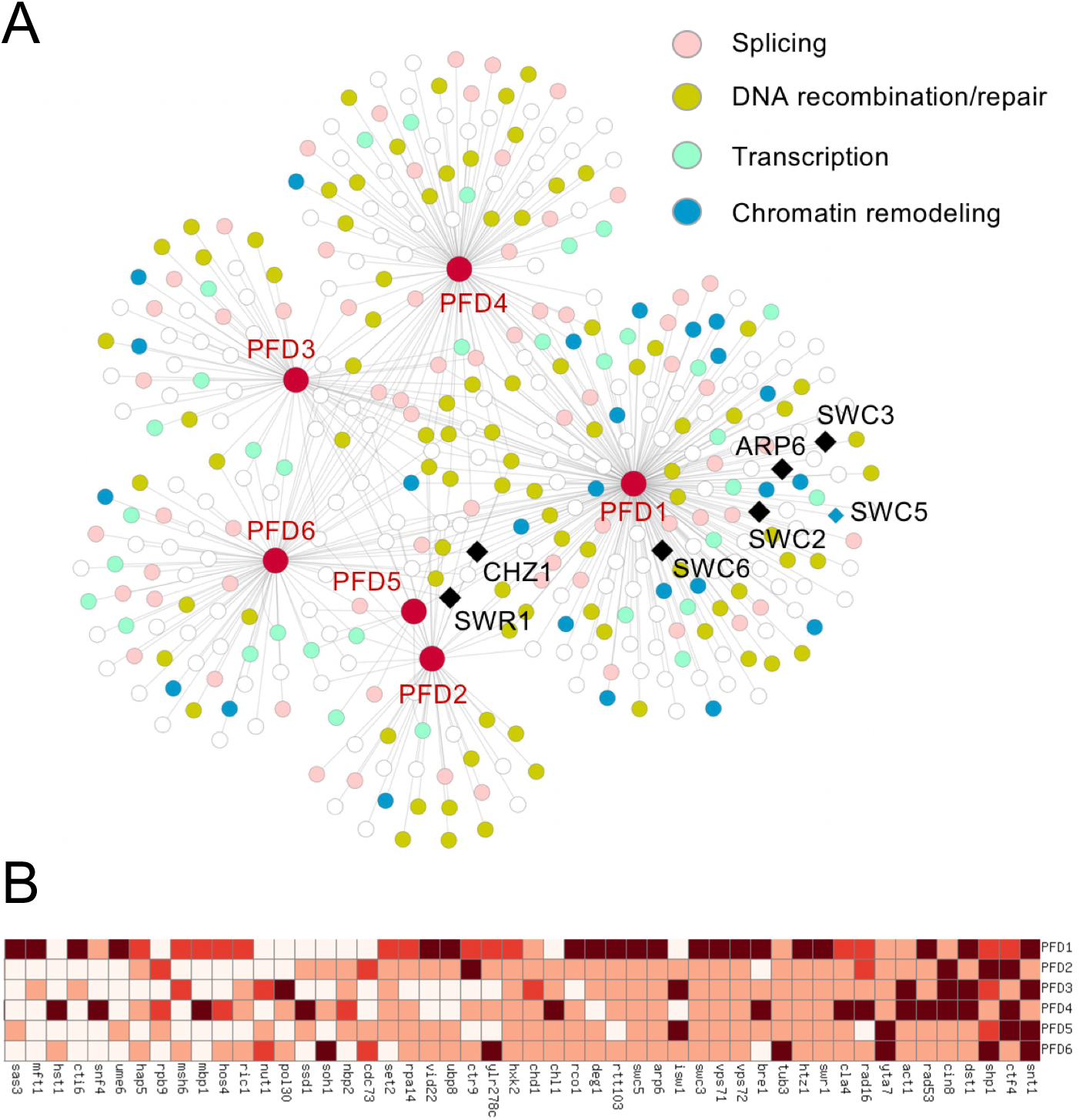
Physical and genetic interactions of yeast PFD subunits. (A) Network representation of physical interactions between PFD subunits and nuclear proteins, highlighting the functions related to DNA biology and subunits of the SWR1 complex. (B) Heatmap representing the top 50 stronger genetic interactions with PFD genes. The darker the color, the stronger the interaction.

**Figure S2.**
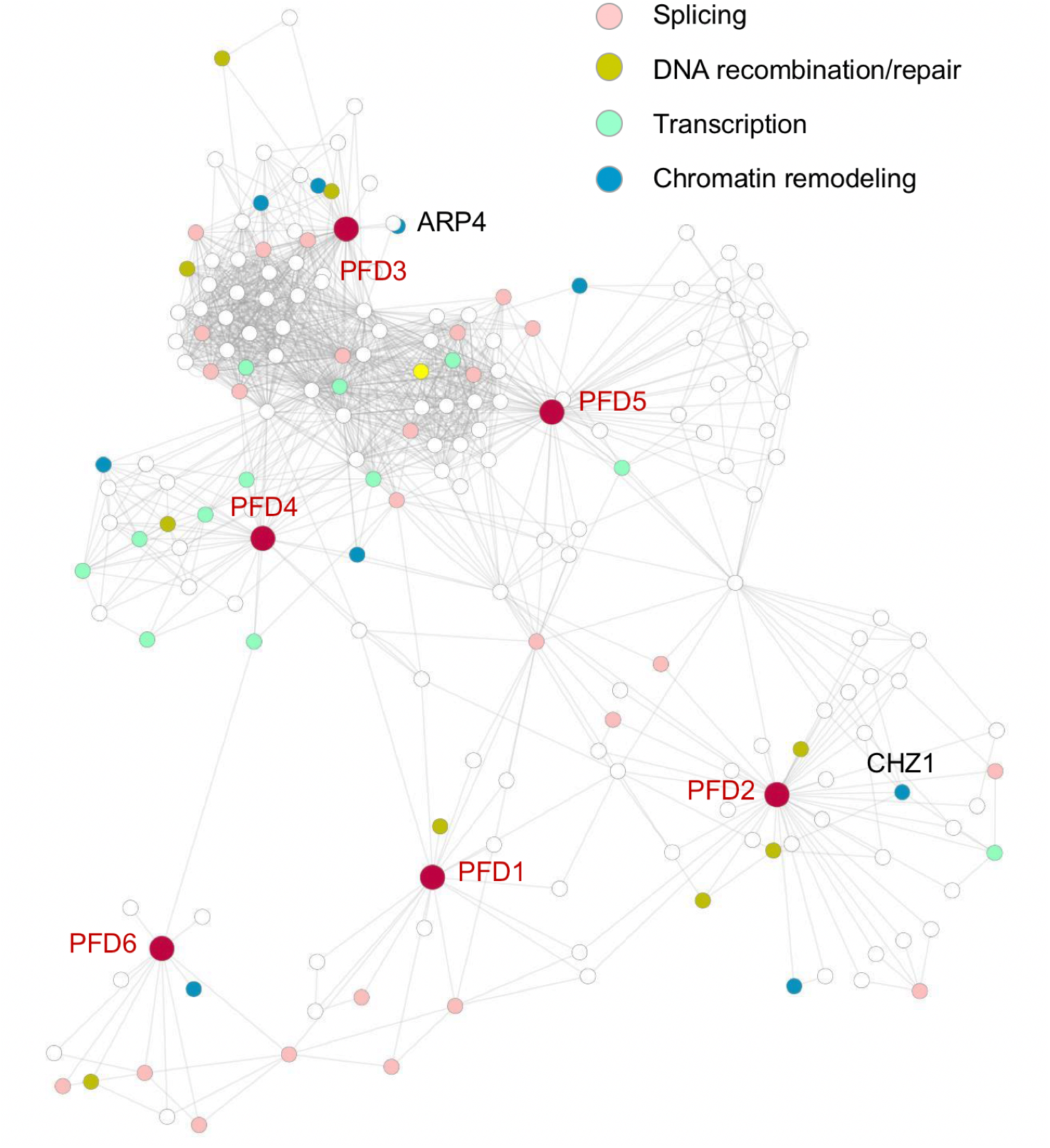
Network representation of the genes significantly coexpressed with *PFD* genes in yeast. Data were extracted from the YeastNet v3 database, and the network was constructed using Cytoscape. CHZ1 and ARP4 are part of the yeast SWR1c.

**Figure S3.**
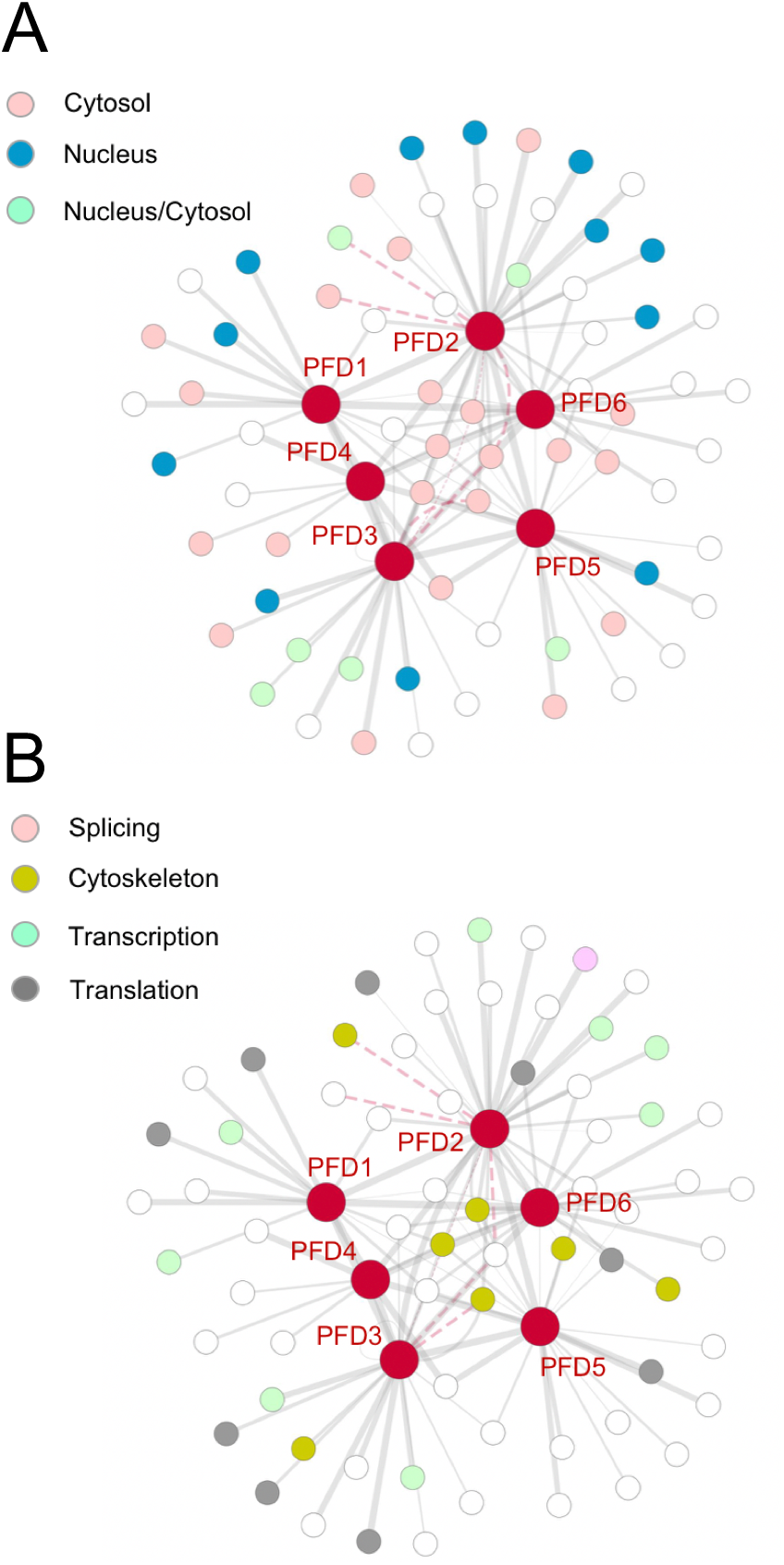
Network representation of physical and genetic interactors of PFD subunits in *D. melanogaster.* Genetic interactions are represented by dashed lines. The width if the edges is directly proportional to the coexpression value. Data were extracted from the InterMine database, and the network was constructed using Cytoscape. (A) Subcellular localization of the interactors. (B) Functional categories of the interactors.

**Figure S4.**
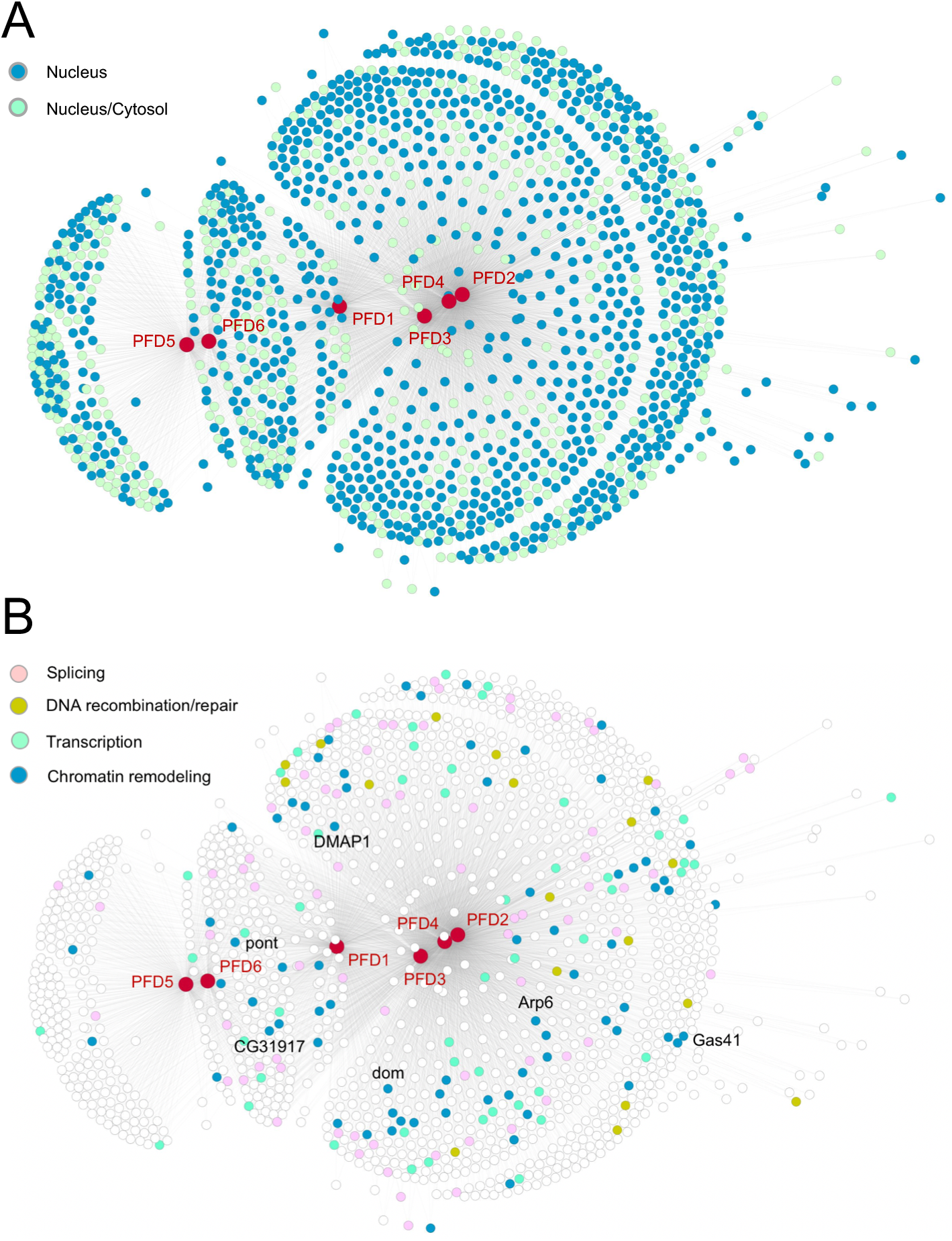
Network representation of the genes significantly coexpressed with *PFD* genes in *D. melanogaster.* Genes encoding subunits of the SWR1 complex are highlighted. (A) Subcellular localization of the interactors. (B) Functional categories of the interactors.

**Figure S5.**
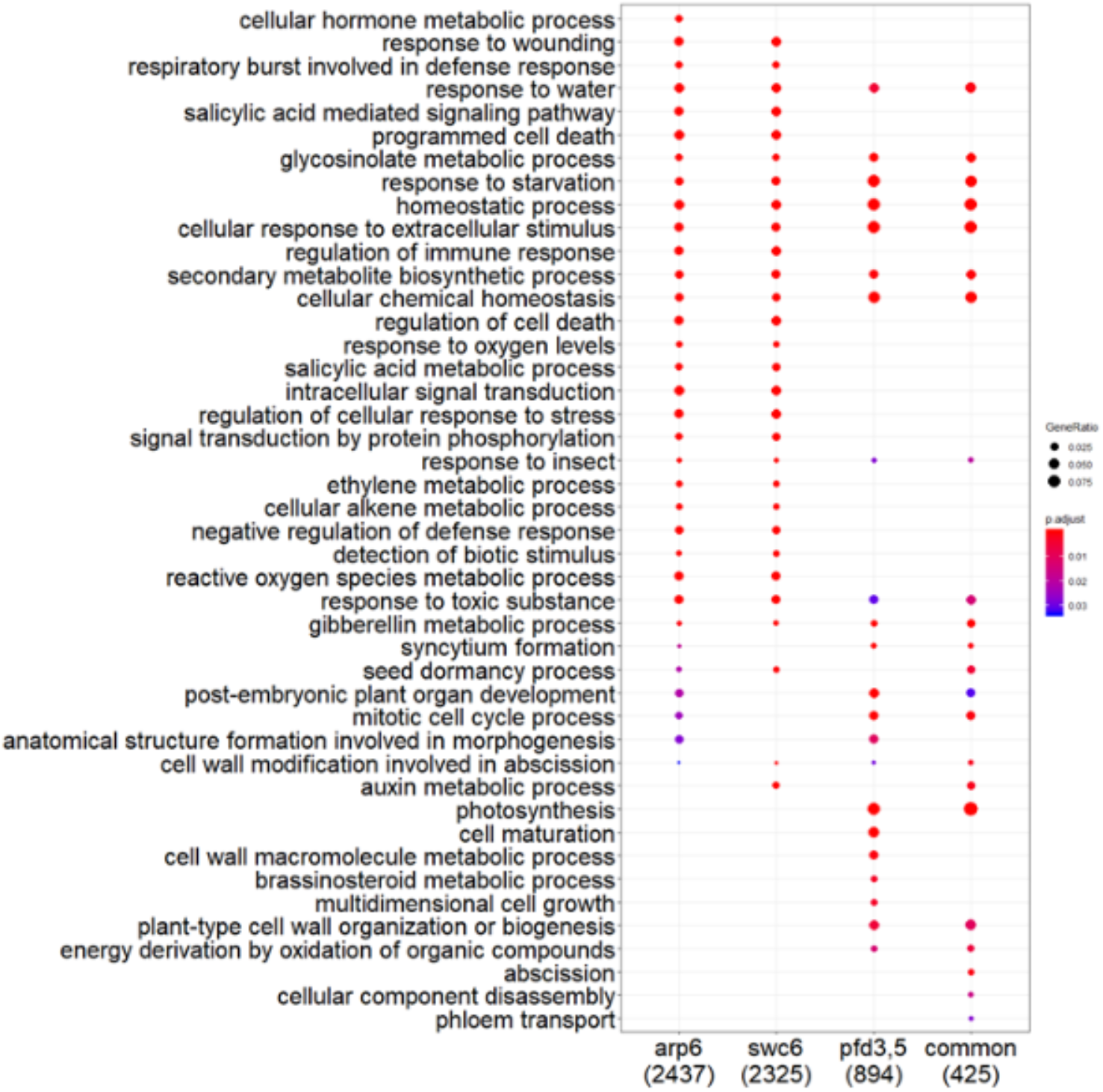
Gene Ontology enrichment analysis of misregulated genes in *arp6, swc6,* and *pfd3,5* mutants. Numbers in brackets indicate the number of misregulated genes in each genotype.

**Figure S6.**
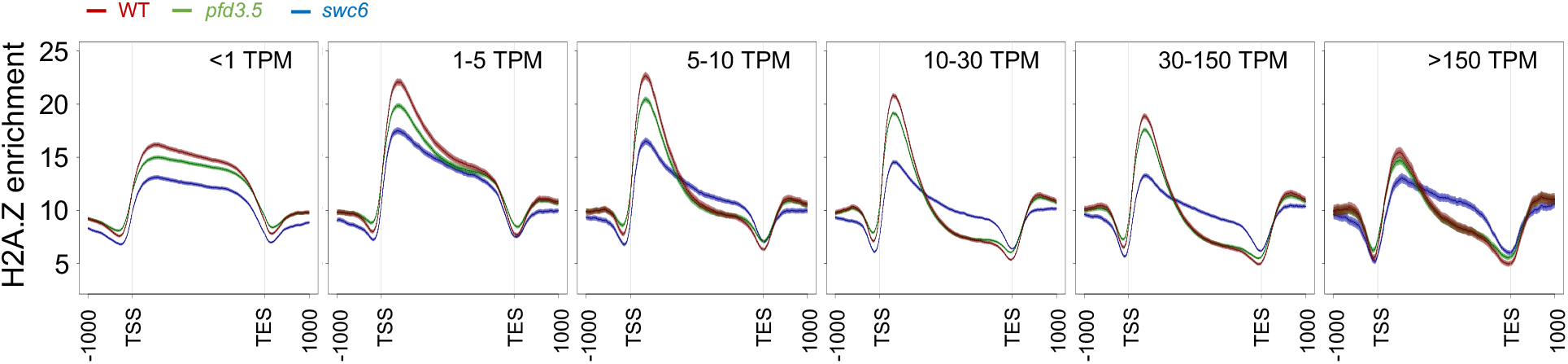
Metagene plots showing H2A.Z levels over different subsets of genes grouped according to their expression levels in the wild-type (indicated in TPM (transcripts per million mapped reads). TSS transcription start site, TES transcription stop site.

**Figure S7.**
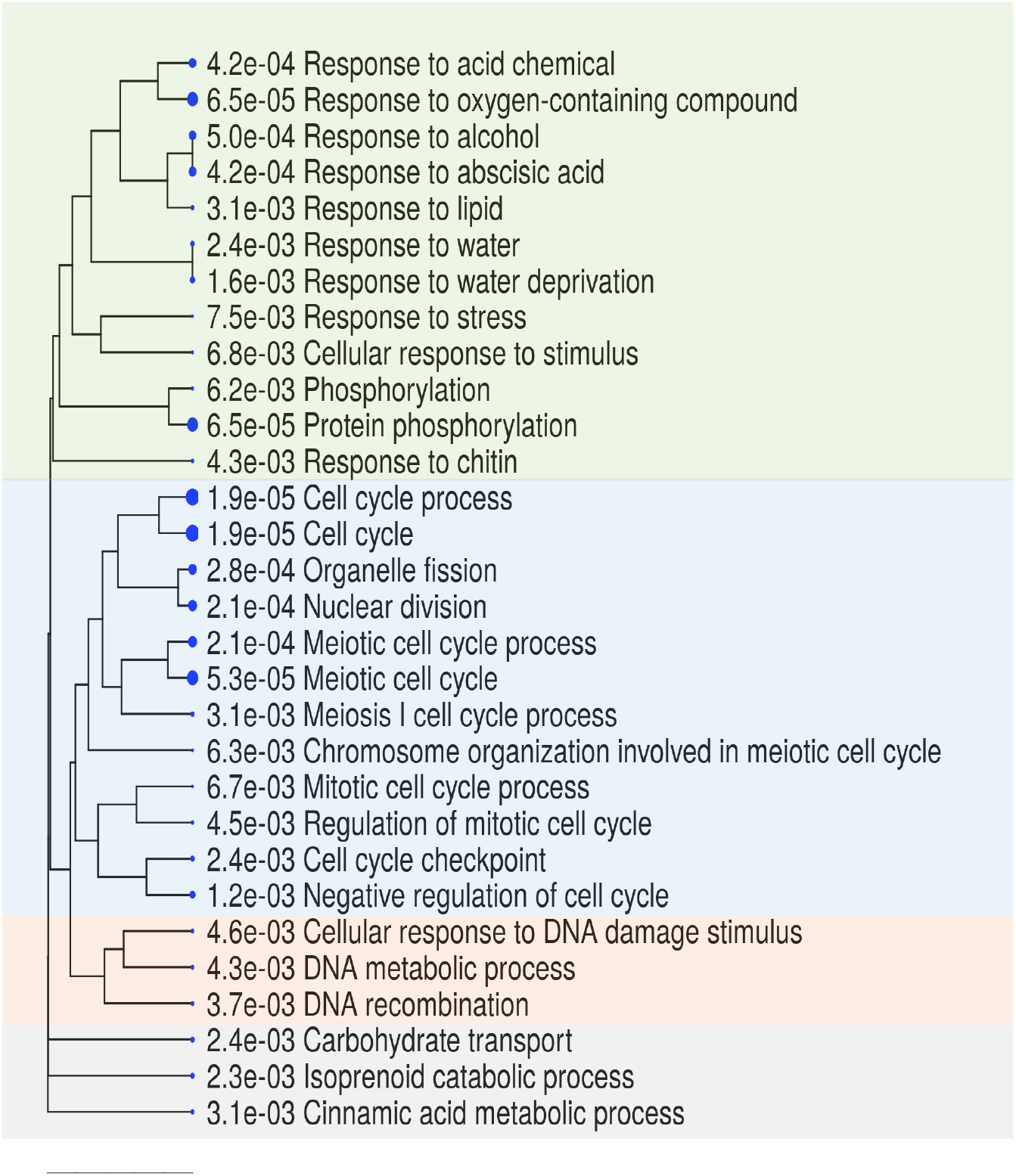
Gene Ontology analysis of genes in which H2A.Z deposition is affected by both mutations.

**Figure S8.**
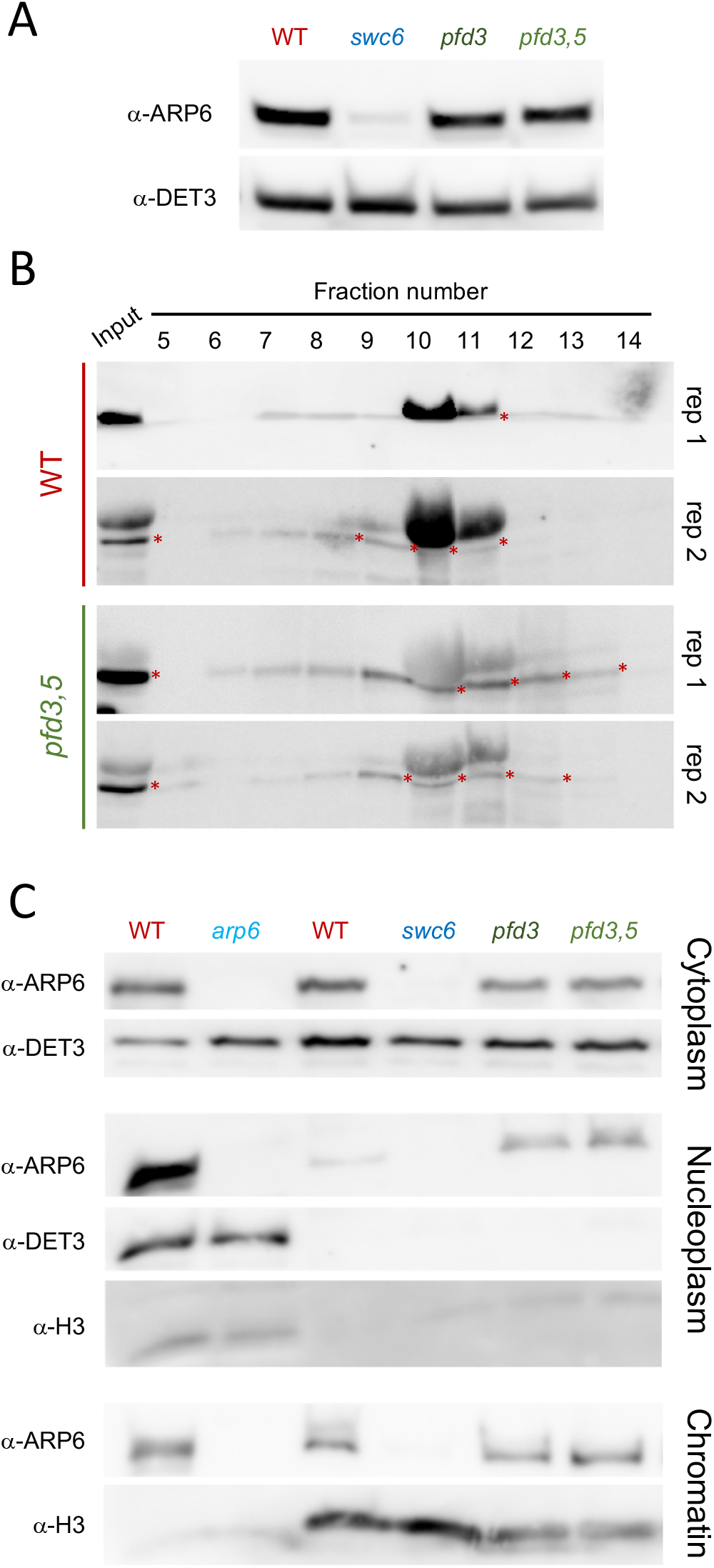
(A) ARP6 protein levels determined by Western analysis. DET3 was used as loading control. (B) Cell fractioning analysis of extracts of the wild-type and *pfd3,5* mutant seedlings. Two replicates are shown. Asterisks mark the band corresponding to ARP6. (C) Cell fractioning shows overaccumulation of the ARP6 subunit in the nucleoplasm of *pfd* mutants. Proteins from the different cellular fractions were subjected to Western analysis. H3 and DET3 proteins were used as controls for chromatin and cytoplasm fractions, respectively.

